# Massive-scale single-cell chromatin accessibility sequencing using combinatorial fluidic indexing

**DOI:** 10.1101/2023.09.17.558155

**Authors:** Xuan Zhang, Alexandre P Marand, Haidong Yan, Robert J Schmitz

## Abstract

Single-cell ATAC-seq has emerged as a powerful approach for revealing candidate *cis*-regulatory elements genome-wide at cell-type resolution. However, current single-cell methods suffer from limited throughput and high costs. Here, we present a novel technique called single-cell combinatorial fluidic indexing ATAC-sequencing (“scifi-ATAC-seq”), which combines a barcoded Tn5 pre-indexing step with droplet-based single-cell ATAC-seq using a widely commercialized microfluidics platform (10X Genomics). With scifi-ATAC-seq, up to 200,000 nuclei across multiple samples in a single emulsion reaction can be indexed, representing a ∼20-fold increase in throughput compared to the standard 10X Genomics workflow.

## Background

ATAC-seq (Assay for Transposase-Accessible Chromatin using sequencing) [1] has emerged as a popular method for chromatin accessibility profiling to unveil genome-wide candidate *cis*-regulatory elements, which govern precise gene expression patterns for specifying distinct cell types or cell states. Recently, ATAC-seq was further modified to profile chromatin accessibility at single-cell resolution (scATAC-seq), which was used to generate several *cis*-regulatory atlases for animals and plants [2-7,35,36]. Most single-cell methods have the capability to generate data for hundreds to thousands of cells/nuclei simultaneously and are categorized as either droplet-based or combinatorial indexed techniques [8-11]. Droplet-based approaches are commonly implemented using microfluidics platforms, such as the commercialized products from companies like 10X Genomics, which offer straightforward handling and consistent data quality compared with combinatorial indexing methods [12].

In droplet-based assays the single-cell suspension is loaded into the microfluidics device at a significantly lower concentration compared to the barcoded beads to minimize the occurrence of cell/nuclei doublets that lead to false cell states. For instance, typically the loading cell/nuclei number is only about 1-10% of the total number of Gel Bead-in-Emulsions (GEMs) from the 10X Genomics Chromium Controller (Fig. 1a), which uses reagents inefficiently, leading to limited throughput and higher costs. To address the inefficiency of droplet-based approaches, a chimeric single-cell strategy has been developed, in which nuclei were pre-indexed prior to overloading of the microfluidics device. With this strategy, the technique known as “dsciATAC-seq” was developed, which combined pre-indexing with a microfluidics system from Bio-Rad, and recovered about 25k nuclei with 100k nuclei loaded in one experiment [13]. More recently, similar approaches have been used for scRNA-seq with the 10X Genomics platform, referred to as scifi-RNA-seq, which recovered around 150k cells upon loading 380k cells, and further significantly increased the throughput [14]. However, there is currently no massive-scale scATAC-seq protocol for use with the 10X Genomics platform, which is most commonly used for scATAC-seq data generation.

**Figure 1.**
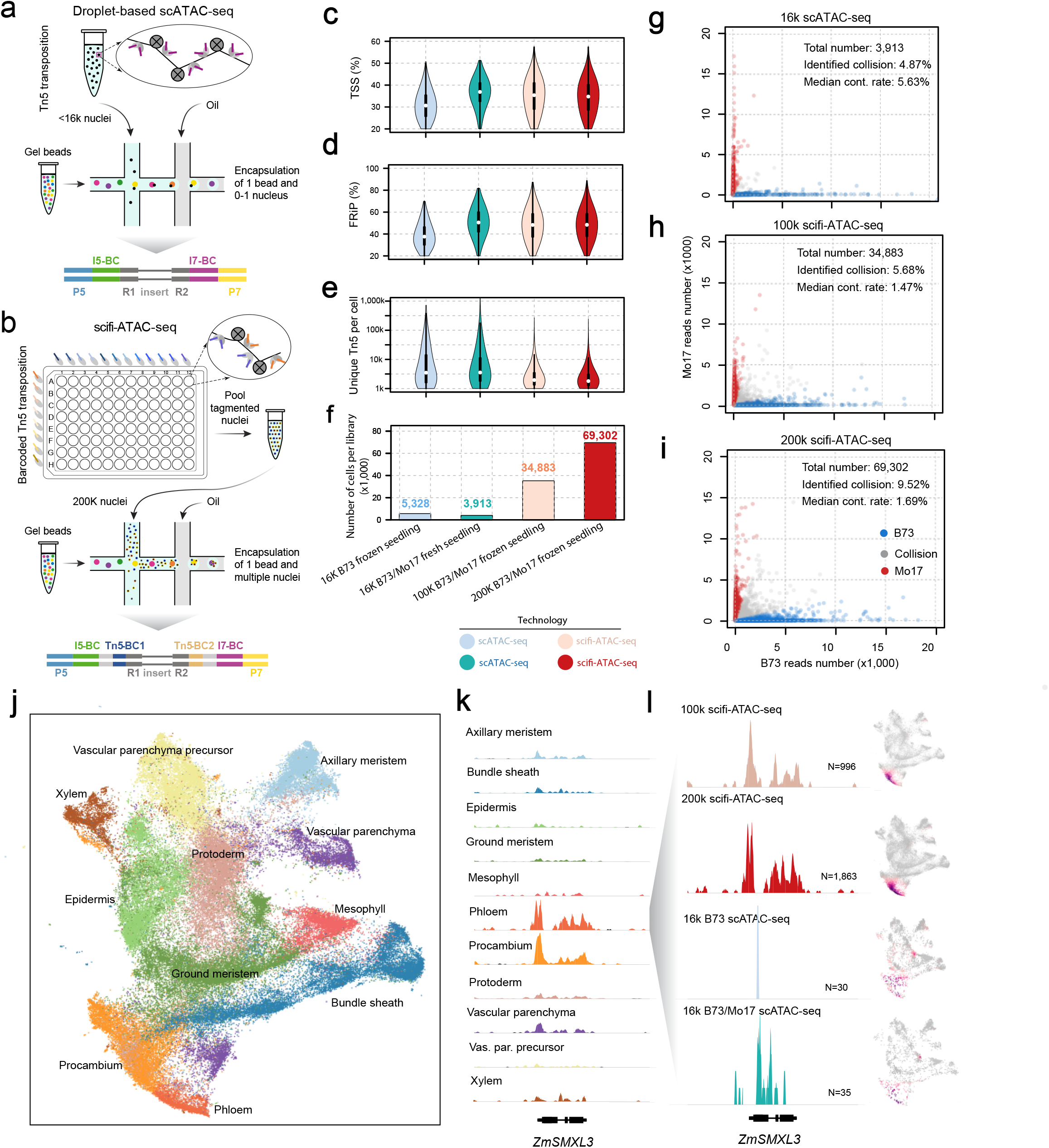
scifi-ATAC-seq combines pre-indexing with droplet-based scATAC-seq. (a) Schematic of regular droplet-based 10X Genomics scATAC-seq experimental workflow. (b) Schematic of scifi-ATAC-seq experimental workflow. (c) Distributions of the proportion of Tn5 integration sites within the promoter regions, encompassing the 2-kb flanking regions around gene transcription start sites (TSSs). (d) Distributions of the proportion of Tn5 integration sites within peaks per nucleus. (e) Distribution of unique Tn5 integration sites per nucleus. (f) Number of nuclei that passed quality control thresholds. (g-i) Scatterplot displaying number of reads per cell classified as B73 or Mo17, color-coded by genotype classification. g, 16k input B73/Mo17 scATAC-seq; h, 100k scifi-ATAC-seq; i, 200k scifi-ATAC-seq. Median Contamination Rate: The median cross-contamination rate, attributed to index hopping, among all predicted singlets. (j) UMAP of all nuclei (n = 98,424). Nuclei are colored by their predicted cell type. (k) Pseudobulk cell type Tn5 integration site coverage around the phloem precursor marker *ZmSMXL3*. Vas. par. precursor: Vascular parenchyma precursor. (l) Pseudobulk cell type Tn5 integration site coverage for and UMAP embeddings overlaid with gene chromatin accessibility around the *ZmSMXL3* gene across all datasets. N: Number of phloem nuclei.

## Results and Discussion

In this study, we present a method for profiling massive-scale single-cell chromatin accessibility sequencing using the 10X Genomics microfluidics system. We refer to this method as single-cell combinatorial fluidic indexing ATAC-sequencing (scifi-ATAC-seq), as it was initially inspired by the scifi-RNA-seq design [14]. In scifi-ATAC-seq, the nuclei are pre-indexed in a 96-well plate with a two-sided barcoded Tn5 (96 distinct barcode combinations), which is based on our previous sci-ATAC-seq design [15]. Then, a standard scATAC-seq library preparation is performed, except that an overloaded number of nuclei are used in the microfluidics system. Approximately 100-200k nuclei per channel instead of the recommended maximum of 15.3k are loaded. Compared to the one-sided barcoded Tn5 [13], the two-sided barcode design offers several advantages: (i) It requires fewer Tn5 adapter oligos for Tn5 barcoding, and readily accommodates scaling up the index complexity when necessary. Only 20 oligos (8 rows x 12 columns) are needed to create 96 unique barcode combinations. (ii) It requires less Tn5 for Tn5 assembly. Only 280 uL of Tn5 (15 uL in 8 rows and 10 uL in 12 columns) is necessary, whereas the one-sided barcode would require over 1,000 uL of Tn5, assuming a minimal assembly volume of 10 uL per well. (iii) The barcode in the s5 end helps to distinguish index hopping reads and reduces index hopping contamination (Fig. S1 a,b).

To evaluate the performance of scifi-ATAC-seq, we generated two scifi-ATAC-seq libraries using 100k and 200k overloaded nuclei from frozen tissue with mixed genotypes (B73 and Mo17) of *Zea mays* (maize) seedling tissue. We compared the scifi-ATAC-seq data with two scATAC-seq libraries generated from the same tissue type using the standard 10X Genomics workflow (referred to as scATAC-seq from this point onward) [4]. All datasets were processed with the same parameters (see Methods). We assessed various quality control metrics, including the proportion of reads around the transcription start site (TSS), fraction of reads in peaks (FRiP), unique Tn5 insertions per cell, fragment size distribution, and organelle DNA contamination (see Fig. 1c-e and Fig. S1c). The scifi-ATAC-seq data showed similar or superior data quality compared to the other datasets. Particularly, scifi-ATAC-seq successfully recovered approximately 35k and 70k clean cells for the 100k and 200k input nuclei, respectively, which represents a 9-fold to 18-fold increase in throughput compared to the scATAC-seq (Fig. 1f). As expected, we observed a higher number of nuclei per droplet when overloading the nuclei, resulting in an average of 1.57 nuclei per droplet for the 100k library and 2.02 nuclei per droplet for the 200k library (Fig. S1 d,f). These data show that scifi-ATAC-seq produces high-quality libraries while increasing the number of nuclei profiled.

Collecting multiple nuclei within a single droplet increases the risk of cell barcode collisions and potential index hopping contamination (Fig. S1a). Next, we compared the barcode collision rate for scATAC-seq and scifi-ATAC-seq with the same approach [4]. The identified barcode collision rate is 4.87% and 5.68% for the 16k scATAC-seq and 100k scifi-ATAC-seq, respectively (Fig. 1g,h). The total barcode collision should be around 10% for both datasets considering the collisions of the same genotype, which is similar with the collision rate reported in other studies [4]. The throughput increased about 9-fold with the 100k scifi-ATAC-seq experiment with a similar barcode collision rate. The identified collision rate was 9.52% for the 200k scifi-ATAC-seq, which is higher than the 16k scATAC-seq dataset. The total recovered nuclei number is further increased to 69,302, which reflects an 18-fold increase in throughput (Fig. 1i). As expected, the total number of the nuclei in the droplet with barcode collisions and the number of Tn5 insertions in nuclei with barcode collisions is significantly higher than non-collision droplets or nuclei (see Fig. S1d-h, q-value < 10e-16). For the droplets containing 1 to 10 nuclei, there is no obvious bias for several quality metrics, such as the proportion of reads around the TSS, FRiP score, unique Tn5 insertions per cell (Fig. S2a-c) whereas there is a noticeable increase in the contamination rate with more cells in the droplet, but it remains at a low level (<5%) even in the droplets with 10 nuclei (Fig S2d). Barcode collisions are typically removed using an array of doublet detection tools [16-19], minimizing the effect of clustering and cell-type identification. Lastly, the cross cell contamination was 1.47% and 1.69% for both scifi-ATAC-seq datasets, which is much lower than the 5.63% observed in the 16k scATAC-seq dataset indicating this could benefit from the double-side barcode design implemented in our method (Fig. S1 a,b). Collectively, these data show that scifi-ATAC-seq has similar barcode collision rates as the standard scATAC-seq workflow, while producing data for significantly greater numbers of nuclei.

To evaluate whether the differences in data quality and cell number could affect clustering, we clustered all the nuclei from the scATAC-seq and scifi-ATAC-seq datasets together with identical parameters using *Socrates* [4] and annotated the clusters using previously reported methods. In total, we identified 14 clusters representing the major cell types in the maize seedling (Fig. 1j, Fig. S3a). All the resulting cell types identified were consistent and found across multiple techniques (Fig. S3b-d). To compare the nuclei quality between scifi-ATAC-seq and scATAC-seq, we randomly selected 100 cells for each cell type or library and checked the distribution of the number of Tn5 integrations for each cell type. We observed that scifi-ATAC-seq has a lower median number of Tn5 integrations (Figure S4a,b). While the lower Tn5 integration in scifi-ATAC-seq does not significantly affect clustering in this study, and the increased numbers of nuclei from scifi-ATAC-seq helps decipher cell heterogeneity (Figure S4 c-e). The ability to identify and characterize rare cell populations in detail is a significant advantage of single-cell technologies, which requires profiling a substantial number of cells within each tissue. Here, we profiled more than 100k nuclei in the seedling tissue with scifi-ATAC-seq, which provides an opportunity to study rarer cell types. In maize seedlings, bundle sheath and mesophyll represent major cell types for photosynthesis and typically represent ∼20% of cell types in seedling tissue, whereas vascular cells, such as phloem, procambium, and xylem, responsible for nutrient transport, are present in much lower proportions (Fig. S5a-d). We observed a similar profile for bundle sheath and mesophyll cells in both scifi-ATAC-seq and the scATAC-seq data (Fig. S5e, f). However, for vascular cells, the resolution of accessible chromatin was difficult to discern in the scATAC-seq data, but were clearly visible in both scifi-ATAC-seq datasets (Fig. 1k, l, Fig. S5g-i). Scifi-ATAC-seq data leads to greater number of profiled nuclei per sample increasing the chances of studying rarer cell populations.

To assess the robustness of scifi-ATAC-seq in profiling single-cell chromatin accessibility across multiple samples within a single reaction, we implemented a multiplexing strategy using maize seedlings from eight samples with seven different genotypes in a 96-well plate (Fig. 2a). With seven different genotypes in this assay, the likelihood of index hopping contamination within the same genotype is expected to be low, as most index hopping occurred between distinct genotypes and can be identified through computational methods. Thus, increasing the number of loaded nuclei, while maintaining a low index hopping contamination rate within the same genotype, can further enhance throughput. To test this, a total of 300k nuclei were used to prepare the scifi-ATAC-seq library. We applied the same nuclei quality control criteria as before and successfully recovered 199,212 high-quality nuclei. We assigned all nuclei with the expected genotypes by matching known Tn5 barcodes and identified 133,524 singlet clean nuclei with a low index hopping contamination rate of 1.93% (Fig. 2b, Fig. S6). As expected, there was high correlation between sequencing throughput and nuclei number for all samples (R=0.98, Fig. S6d). The estimated barcode collision rate ranged from 1.0% to 7.6% (Fig. 2b, Fig. S6e-l, Methods). To further validate the biological relevance and quality of the data, we conducted a Spearman correlation analysis with chromatin accessibility among the eight libraries (Fig 2c). Our findings reveal that the clustering of different genotypes aligns well with maize genetic divergence, notably grouping all genotypes from non-stiff stalk (NSS) together. These results demonstrate that the throughput was further enhanced with scifi-ATAC-seq by approximately 27-fold compared to the scATAC-seq method, while maintaining a similar barcode collision rate and data quality.

**Figure 2.**
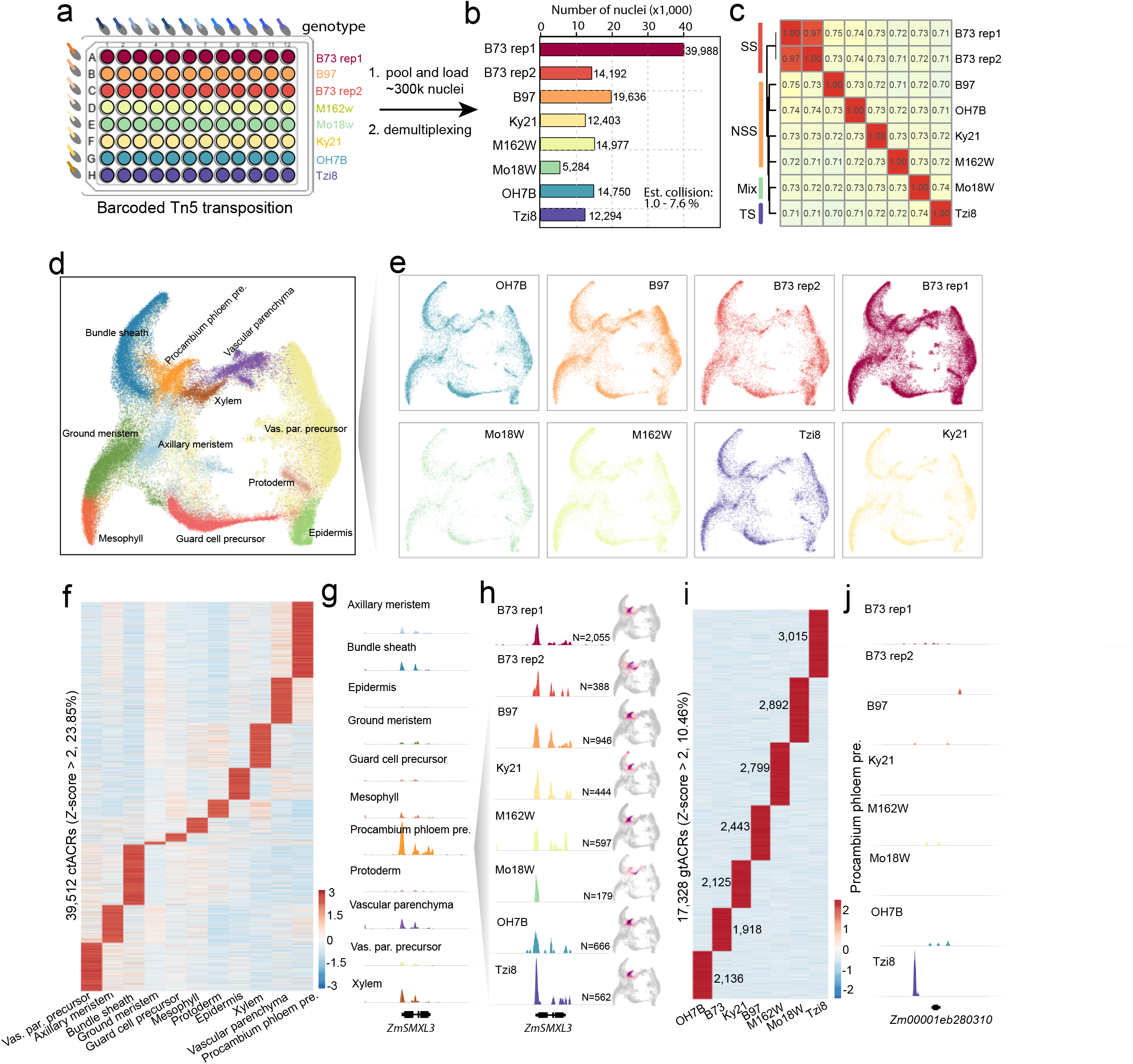
Multiplexing eight samples with scifi-ATAC-seq. (a) Well assignment showing the multiplexing of primary samples and genotypes. (b) Number of profiled nuclei for each genotype. Est. Collision: Estimated cell barcode collisions within the same genotype. (c) Spearman Correlation heatmap among the 8 libraries.(SS, stiff stalk; NSS, non-stiff stalk; TS, tropical/subtropical; Mixed, mixed tropical-temperate) (d) UMAP of all nuclei (n = 124,656). Nuclei are colored by their predicted cell type. Vas. par. precursor: Vascular parenchyma precursor, Procambium phloem pre.: Procambium phloem precursor. (e) UMAP of all nuclei across the eight samples. (f) Chromatin accessibility of cell-type-specific accessible chromatin regions (ctACRs, *Z*-score > 2) among all cell types. (g) Pseudobulk cell type Tn5 integration site coverage around the phloem precursor marker *ZmSMXL3*. (h) Pseudobulk cell type Tn5 integration site coverage and UMAP embeddings overlaid with gene chromatin accessibility around the *ZmSMXL3*. N: Nuclei number of procambium phloem precursor. (i) Chromatin accessibility of genotype specific accessible chromatin regions (gtACRs) among seven genotypes. The number of gtACRs was labled. (j) Pseudobulk cell type Tn5 integration site coverage with gene chromatin accessibility around the *Zm00001eb280310* across all samples in procambium phloem precursor cells.

Subsequently, we clustered all the singlet nuclei from this assay and annotated the clusters using the same method as before. We identified all the cell types observed in the previous dataset and observed that there was no significant bias in cell-type proportions among the genotypes (Fig. 2d, e, Fig. S7d). With this dataset, we identified 165,666 accessible chromatin regions, with approximately 23.85% showing cell-type-specific patterns and 10.46% showing genotype-specific patterns (Fig. 2f, i). More specifically, vascular cells (procambium, phloem, and xylem) were consistently identified in all samples (Fig. 2g, Fig. S8a-c), and genotype-specific accessible chromatin regions at cell-type resolution were discernible (Fig. 2j, Fig. S8d). For the tropical line Tzi8, the gtACR-associated genes were most enriched in photomorphogenesis (p-value < 1e-4) and regulation of response to red or far-red light (p-value < 1e-3). These findings may reflect its distinct adaptation response to day length or light intensity (Table S8,9).

## Conclusions

In summary, these results collectively show that scifi-ATAC-seq provides a robust, efficient, and flexible approach for massive-scale single-cell chromatin accessibility profiling using the widely available 10X Genomics Chromium systems. We anticipate that this method will facilitate the utilization of ATAC-seq to identify candidate *cis*-regulatory elements at cell-type resolution in greater numbers of tissues, time point, genotypes and facilitate the study of rarer cell types.

## Methods

### Plant material and growth conditions

The maize kernels used in this study were obtained from the USDA National Plant Germplasm System (https://npgsweb.ars-grin.gov). The seedlings were grown in Sungro Horticulture professional growing mix (Sungro Horticulture Canada) under controlled conditions. The soil was saturated with tap water and the seedlings were exposed to a mixture of 4,100 K (Sylvania Supersaver Cool White Delux F34CWX/SS, 34 W) and 3,000 K (GE Ecolux with starcoat, F40CX30ECO, 40 W) light, with a photoperiod of 16 hours of light and 8 hours of darkness. The temperature was maintained at approximately 25°C during the light hours, and the relative humidity was approximately 54%. The above-ground seedling tissues were harvested between 8 and 9 AM, six days after sowing. Flash-frozen seedling tissue was used to generate scifi-ATAC-seq libraries for B73 and Mo17 mixed genotypes. And fresh seedling tissue were used for scifi-ATAC-seq library with seven different maize genotypes.

### scifi-ATAC-seq protocol

A detailed step-by-step sci-ATAC-seq protocol with lists of necessary reagents and equipment are included in the Supplementary Methods. The Tn5 expression and purification steps were carried out according to the method described by Tu et al. (2020) [20], and the plasmids were obtained from Addgene (accession number 127916).

### Assembly of indexed Tn5 transposase complexes

To generate indexed Tn5 transposase complexes, we modified the Tn5-ME-A and Tn5-ME-B by adding a 5-nucleotide barcode (with 12 distinct barcodes for Tn5-ME-A and 8 distinct barcodes for Tn5-ME-B; see Supplementary Methods Supplementary Table 1 for barcode sequences). Each indexed oligonucleotide was annealed to a 19-bp complementary mosaic-end oligonucleotide (Tn5-ME-rev, 5′ phosphorylated, Supplementary Methods). The annealing reaction was carried out by mixing the oligonucleotides at a 1:1 molar ratio (Tn5-ME-A or Tn5-ME-B: Tn5-ME-rev) at a final concentration of 100 μM. The mixture was heated to 95 °C, cooled gradually to 20 °C at a rate of -1 °C per minute, and then held at 20 °C. The annealed oligonucleotides were then mixed at a 1:1 molar ratio (Tn5-ME-A: Tn5-ME-B). To assemble the transposase, 10 μL of Tn5 transposase was added to 0.143 μL of the annealed adapter mixture, mixed well by pipetting slowly, incubated at room temperature for 60 minutes and then storing it at -20°C until the tagmentation reactions were performed [21].

### Nuclei isolation with quick purification

Approximately 3-4 maize seedlings were chopped on ice for about 2 minutes in 600 μL of pre-chilled Nuclei Isolation Buffer (NIB cutting, 10 mM MES-KOH pH 5.4, 10 mM NaCl, 250 mM sucrose, 0.1 mM spermine, 0.5 mM spermidine, 1 mM DTT, 1% BSA, 0.5% TritonX-100), which was modified from the original buffer composition [22]. After chopping, the total mixture was filtered with a 40-μm cell strainer and then centrifuged at 500 rcf for 5 minutes at 4 °C. The supernatant was carefully removed, and the pellet was resuspended in 500 μL of NIB wash buffer (10 mM MES-KOH, pH 5.4, 10 mM NaCl, 250 mM sucrose, 0.1 mM spermine, 0.5 mM spermidine, 1 mM DTT, and 1% BSA). The sample was filtered with a 20-μm cell strainer and then carefully loaded onto the surface of 1 mL 35% Percoll buffer (made by mixing 35% Percoll and 65% NIB wash buffer) in a 1.5-mL centrifuge tube. The nuclei were centrifuged at 500 rcf for 10 minutes at 4 °C. After centrifugation, the supernatant was carefully removed, and the pellets were washed once in 100 μL TAPS buffer (25 mM TAPS-NaOH, pH 8.0, and 12.5 mM MgCl_2_) and then resuspended in 30 μL of 2.5x TAPS buffer. About 5 μL of nuclei were diluted 10 times and stained with DAPI (Sigma Cat. D9542). The nuclei quality and density were evaluated with a hemocytometer under a microscope. Finally, after nuclei for both genotypes were isolated separately, equal nuclei number of B73 and Mo17 were mixed together and the nuclei density was adjusted to 0.5k∼1k/μL with TAPS buffer.

### Indexed Tn5 tagmentation and pooling

To generate a combination of 96 indexed transposases, 1.5 μL of Tn5-ME-A with 12 distinct barcodes were dispensed by rows, and 1.5 μL of Tn5-ME-B with 8 distinct barcodes were dispensed by columns in a 96-well plate. Each well had a unique combination of A and B indexed Tn5. To each well, 10 μL of nuclei in TAPS buffer with 0.1% Tween 20 and 0.01% digitonin was added, and the plate was sealed. The tagmentation reaction was carried out for 60 minutes at 37 °C. The reaction was stopped by adding 12 μL of stop buffer (10 mM Tris-HCl pH 7.8, 20 mM EDTA, pH 8.0, 2% BSA) supplemented with ethylenediaminetetraacetic acid (EDTA) to quench the Mg2+. All nuclei were transferred to a reservoir and then divided into two 1.5 mL centrifuge tubes. The nuclei were pelleted, resuspended in 200 μL diluted nuclei buffer (DNB, 10x Genomics Cat#2000207), filtered with a 40 um strainer, pooled into one PCR tube, and centrifuged at 500 rcf for 2 min at 4°C. After centrifugation, the supernatant was carefully removed, and the nuclei (approximately 3 μL) were resuspended in 5 μL of DNB and 7 μL ATAC buffer B (10x Genomics Cat#2000193).

### Library preparation and sequencing

scATAC-seq libraries were prepared using the Chromium scATAC v1.1 (Next GEM) kit from 10xGenomics, following the manufacturer’s instructions. (10xGenomics, CG000209_Chromium_NextGEM_SingleCell_ATAC_ReagentKits_v1.1_UserGuide_RevE). The leftover nuclei after loading to ChIP-H were diluted, stained with DAPI, and nuclei quality and density was evaluated with a hemocytometer under a microscope. The final libraries were sequenced using an Illumina NovaSeq 6000 S4 in the dual-index mode using custom sequencing primer sets (Supplementary Methods). To balance the nucleotide distribution at the beginning of the forward and reverse reads, the proportion of the scifi-ATAC-seq library in a lane should be less than 50% or extra spike-in library (e.g., PhiX control from Illumina) should be added to the lane. The libraries were sequenced to an average depth of 7,617 read pairs per cell, with an average unique reads rate at 62.3%. The scATAC-seq libraries were sequenced about 51.7k reads pairs per cell [4].

### Raw reads processing and alignment

During the preprocessing of all single-cell ATAC-seq data, the 16-bp i5 beads barcode was added to the read names of the paired-end reads using the extract function from UMItools v.1.01 [23]. The customization parameter ‘--bc-pattern=NNNNNNNNNNNNNNNN’ was used for this process. Moreover, for scifi-ATAC-seq data, the inline Tn5 barcode was demultiplexed and subsequently appended to the read names using cutadapt v3.4 [24]. Next, the processed reads were aligned to the *Zea mays* reference genome v5 [25] using BWA-MEM v0.7.17 [26]. To obtain high-quality, properly paired, and unique alignments, the view function from samtools v1.9 [27] was applied with the parameters ‘-q 10 -f 3.’ Additionally, reads with XA tags were filtered out. Subsequently, the cell barcodes were included in the alignments using the CB tag and BC tag for the 10X Genomics scATAC-seq and scifiATAC-seq datasets, respectively. To eliminate duplicate reads, Picard Tools v.2.21.6 (http://broadinstitute.github.io/picard/) was employed while considering the cell barcode. Finally, the alignments were converted to single base-pair Tn5 integration sites in a BED format by adjusting the start coordinates of the forward and reverse strands by +4 and -5, respectively. Only unique Tn5 insertion sites within a cell were retained for downstream analysis.

### Nuclei calling and quality control

The R package *Socrates* [4]was utilized for nuclei identification and quality control. In summary, the BED file containing single base-pair Tn5 integration sites was imported into *Socrates* along with the *Zea mays* v5 GFF gene annotation and the genome index file. The scaffolds (‘scaf_23’, ‘scaf_34’, and ‘scaf_36’) were considered as organelle genomes. To identify bulk-scale ACRs (Accessible Chromatin Regions) in *Socrates*, the callACRs function was employed with the following parameters: genome size = 8.5e8, shift = -75, extsize = 150, and FDR = 0.1. This step allowed us to estimate the fraction of Tn5 integration sites located within ACRs for each nucleus. Metadata for each nucleus were collected using the buildMetaData function, using a TSS (Transcription Start Site) window size of 2 kb (tss.window = 2000). Subsequently, sparse matrices were generated with the generateMatrix function, using a window size of 500. High-quality nuclei were identified based on the following criteria: a minimum of 1,000 Tn5 insertion sites per nucleus, at least 20% of Tn5 insertions within 2 kb of TSSs, and at least 20% of Tn5 insertions within ACRs across all datasets. Additionally, a maximum of 30% of Tn5 insertions in organelle genomes was allowed.

### Cell barcode collision detection

The term ‘cell barcode’ refers to the barcode used to determine cell identity. In the standard scATAC-seq, the cell barcode corresponds exclusively to the barcode derived from hydrolyzed GEM beads following microfluidic partitioning. However, in scifi-ATAC-seq, the cell barcode encompasses both the beads barcode and the Tn5 barcode. Cell barcode collision occurs when more than one cell shares the same cell barcode. In traditional droplet-based assays, cell barcode collision occurs if multiple cells enter one droplet. In scifi-ATAC-seq, cell barcode collision happens when multiple cells occupy the same droplet, and simultaneously, they share identical Tn5 barcodes. Cell barcode collisions were identified using a previously described approach [4]. Specifically, the known genotype data were obtained from Panzea[28] and lifted over to v5 genome [25] coordinates using CrossMap (v0.5.1) [29]. Only homozygous biallelic SNPs were retained for further analysis. The Souporcell (git version 6872d88) pipeline [30] was used to count the number of reads for each genotype, using known common variants and specific non-default parameters (--min_alt 50, --min_ref 50, --max_loci 25000000, -- skip_remap TRUE). In the dataset comprising a mixture of two genotypes (B73 and Mo17), genotype identification was performed by modeling allele counts as a binomial distribution, accounting for a conservative sequencing error rate of 0.05. Posterior probabilities were estimated via Bayes theorem to assign the genotype or identify cell barcode collisions (i.e., mixtures of genotypes) with the highest probability. A minimum threshold of 50 reads covering common variants within a cell was used to confidently assign the genotype. The total SNP number between genotypes is around 1.6 million and the SNP rate is about 0.70/kb.The index hopping contamination was calculated as the proportion of reads that did not match the assigned genotype among all reads covering the biallelic SNPs. In the dataset featuring a mixture of two genotypes, cell doublets can originate from either two cells of the same genotype (A x A or B x B) or cells from different genotypes (A x B or B x A). However, the observed collisions consist entirely of cell doublets from different genotypes, representing only approximately half of the collision events that actually take place in the experiment. Therefore, there should be an equal proportion of doublets mixed with nuclei of the same genotype. For the seven-genotype-mixed scifi-ATAC-seq data, expected genotypes were assigned by matching known Tn5 barcodes. Any reads that did not match the expected genotype were considered as belonging to another genotype. The same genotype calling approach described above was then used to assign nuclei to their expected genotype, identify mixtures of genotypes resulting from index hopping contamination. Given the varying cell numbers per well, we used a modeling-based approach to estimate the barcode collision rate by calculating the probability of obtaining any two cells from the same well in a four-nuclei droplet (mean nuclei number per droplet is 3.4, Fig S6c).

### Integrated clustering analysis

The integrated clustering analysis of the four datasets, combining scifi-ATAC-seq and 10X Genomics scATAC-seq data, was performed using the R package *Socrates* [4]. In brief, firstly, the ACRs were identified by treating each library as a traditional bulk ATAC-seq library with function callACRs (genomesize=8.5e8, shift= -75, extsize=150, fdr=0.1). Then a binary nucleus x ACR matrix was generated with the function generateMatrix (peaks=T). The ACRs that were accessible in less than 0.15% of all nuclei, and nuclei with less than 100 accessible ACRs were removed. Then the filtered nucleus x ACR matrix were normalized with the term-frequency inverse-document-frequency (TF-IDF) algorithm(doL2=T). The dimensionality of the normalized accessibility scores was reduced using the function reduceDims (method=“SVD”, n.pcs = 25, cor.max =0.5). The reduced embedding was visualized as a UMAP embedding using projectUMAP (k.near = 15). Approximately 5% of potential cell doublets were identified and filtered by performing a modified version of the *Socrates* workflow on each library separately with the function detectDoublets and filterDoublets (removeDoublets = T). To address genotype and batch effects, we used the R package Harmony with non-default parameters (do_pca=F, vars_use=c(“library”, “genotype”), tau=c(5), lambda=c(0.1, 0.1), nclust=50, max.iter.cluster=100, max.iter.harmony=30). The dimensionality of the nuclei embedding was further reduced with Uniform Manifold Approximation Projection (UMAP) via the R implementation of umap (n_neighbors=30, metric = “cosine”, a = 1.95, b = 0.75, ret_model=T). Finally, the nuclei were clustered with function callClusters (res=0.4, k.near = 30, cl.method = 4, m.clust = 100).

A similar clustering process was applied to the 7-genotype-mixed scifi-ATAC-seq dataset with minor modifications. Specifically, we removed only the genotype effect using Harmony, and the final clusters were identified at a resolution of 0.5.

### Cell-type annotation

To assign cell types for each cluster, we used a combination of marker gene-based annotation and gene set enrichment analysis. Initially, we compiled a list of known cell-type-specific marker genes for maize seedlings through an extensive literature review, primarily referring to Marrand et al., 2021 (Supplementary Table 5) [4]. Firstly, the gene chromatin accessibility score was calculated using the Tn5 integration number in the gene body, 500-bp upstream and 100-bp downstream region, then the raw counts were normalized with cpm function in edgeR. The Z-score was calculated for each marker gene across all cell types with scale function in R, and key cell types were assigned based on the most enriched marker genes with highest Z-score. Ambiguous clusters displaying similar patterns to key cell types were assigned to the same cell type as the key cell types, reflecting potential variations in cell states within a cell type. For gene set enrichment analysis, we used the R package fgsea [31], following a methodology described previously[4]. Firstly, we constructed a reference panel by uniformly sampling nuclei from each cluster, with the total number of reference nuclei set to the average number of nuclei per cluster. Subsequently, we aggregated the read counts across nuclei in each cluster for each gene and identified the differential accessibility profiles for all genes between each cluster and the reference panel using the R package edgeR. For each cluster, we generated a gene list sorted in decreasing order of the log2 fold-change value compared to the reference panel and utilized this list for gene set enrichment analysis. We excluded GO terms with gene sets comprising less than 10 or greater than 600 genes from the analysis and GO terms were considered significantly enriched at an FDR < 0.05 with 10,000 permutations. The cell type annotation was additionally validated by identifying the top enriched GO terms that align with the expected cell type functions (Supplementary Table 6,7).

### ACR identification

Following cell clustering and annotation, ACRs were further identified using all Tn5 integration sites for each cell type and genotype with running MACS2 [32] with non-default parameters: --extsize 150 --shift -75 --nomodel --keep-dup all --qvalue 0.05. Then the cell type based ACRs for each genotype were further redefined as 500-bp windows centered on the ACR coverage summit. To consolidate information across all clusters and genotypes, we concatenated all ACRs into a unified master list using a custom script, as previously described by Marrand et al., 2021 [4], calculated the ACR chromatin accessibility score based on the Tn5 integration count within the ACR region and then normalized it using the ‘cpm’ function in edgeR [33]. ACRs with less than 3 cpm in all cell types and genotypes were removed for downstream analysis.

## Supporting information

Supplementary Method scifi-ATAC-seq protocol

## Declarations

### Ethics approval and consent to participate

Not applicable

### Consent for publication

Not applicable

### Availability of data and materials

All data supporting the results of this study are available within the article and supplementary information files. The scifi-ATAC-seq data generated in this study have been deposited in the National Center for Biotechnology Information Short Reads Archive (PRJNA996051).

The code used for data analysis is available at https://github.com/schmitzlab/scifi-ATAC-seq/

### Author contributions

R.J.S. and X.Z. designed the research. X.Z. performed the experiments. A.P.M., H.Y., X.Z., and R.J.S. analyzed the data. R.J.S. and X.Z. wrote the manuscript. The authors read and approved the final manuscript.

## Acknowledgments

We would like to acknowledge Dr. Pedro Augusto Braga Dos Reis, Dr. Yinxin Dong, Yangyang Xu, for HS-Tn5 transposase purification; the GACRC for providing valuable assistance; and the Duke University School of Medicine for the use of the Sequencing and Genomic Technologies Shared Resource, which provided illumina sequencing service.

---

**Figure S1.**
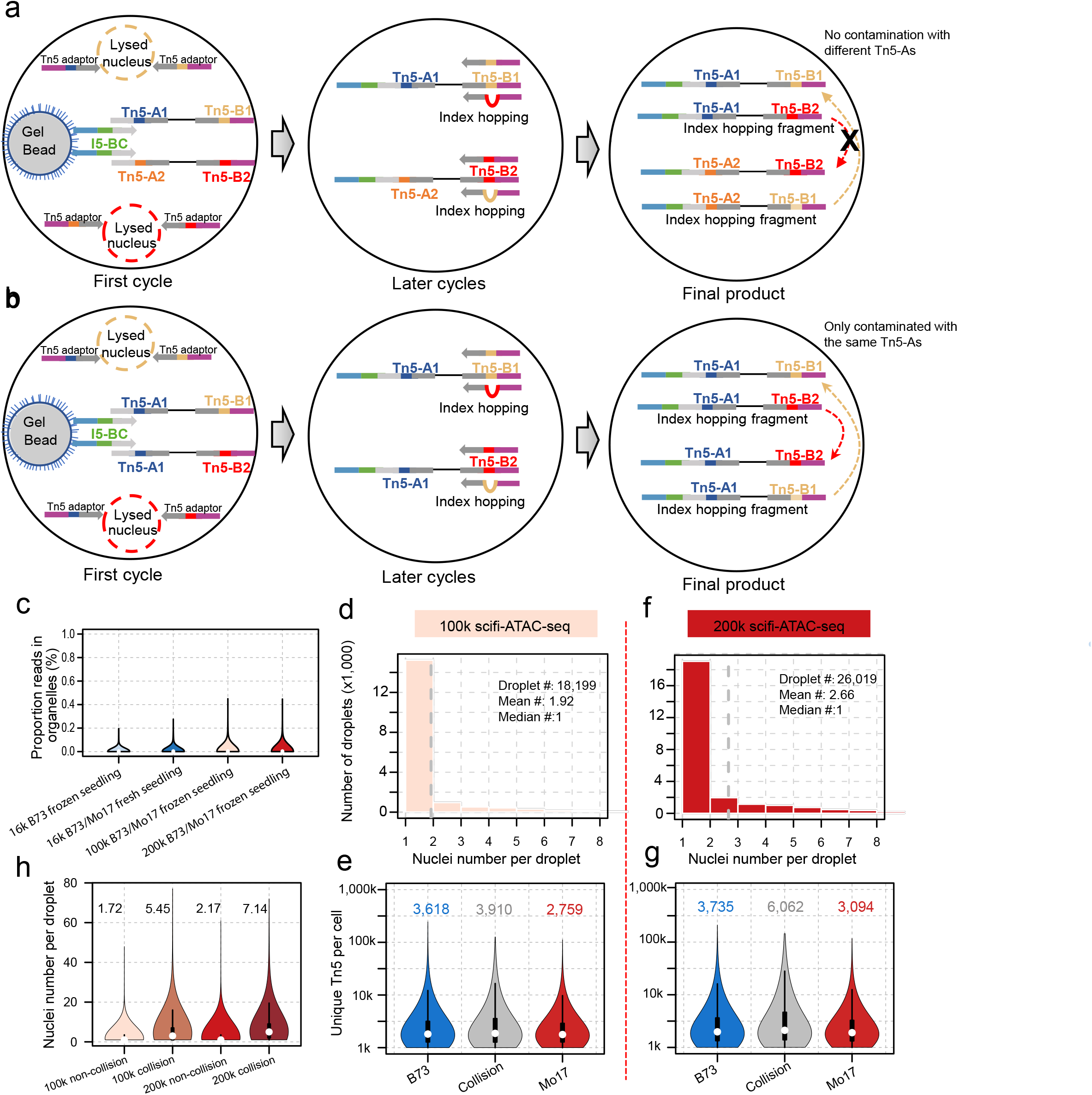
Evaluation and quality control of scifi-ATAC-seq, related to Figure1. (a-b) Illustration of index hopping occurring during linear amplification in 10X Genomics Gel Beads-in-emulsion (GEMs) for scifi-ATAC-seq with (a) distinct Tn5-A barcoded nuclei, ensuring no contamination between nuclei within the droplet; or (b) same Tn5 barcoded nuclei, resulting in contamination between nuclei within the droplet. (c) Distributions of the proportion of Tn5 integration sites within organelle genomes. (d) Number of nuclei per droplet in 100k scifi-ATAC-seq. (e) Distributions of unique Tn5 integration sites across predicted droplet type in 100k scifi-ATAC-seq. (f) Number of nuclei per droplet in 200k scifi-ATAC-seq. (g) Distributions of unique Tn5 integration sites across predicted droplet type in 200k scifi-ATAC-seq. (h) Distributions of nuclei number per droplet for non-collision droplet and collision droplet in 100k and 200k scifi-ATAC-seq. The mean nuclei number labeled in each plot.

**Figure S2.**
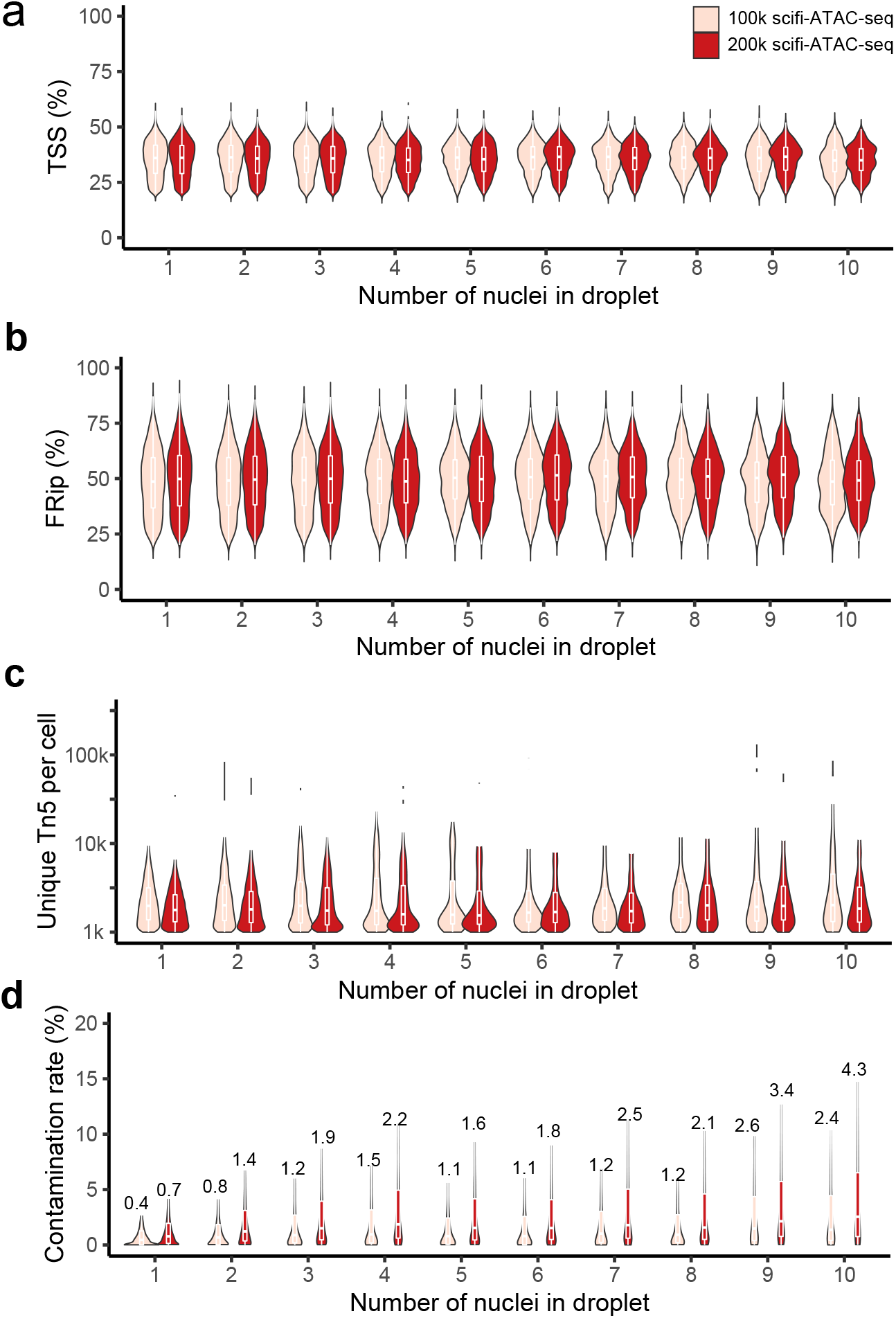
Evaluation and quality control of scifi-ATAC-seq, related to Figure 1. (a) Distributions of proportion of Tn5 integration sites within the promoter regions for droplets containing different nuclei numbers. (b) Distributions of proportion of Tn5 integration sites within peaks for droplets containing different nuclei numbers. (c) Distributions of unique Tn5 integration sites for droplets containing different nuclei numbers. (d) Distributions of index hopping conamination rate for droplets containing different nuclei numbers. The median contamination rate showed in each plot.

**Figure S3.**
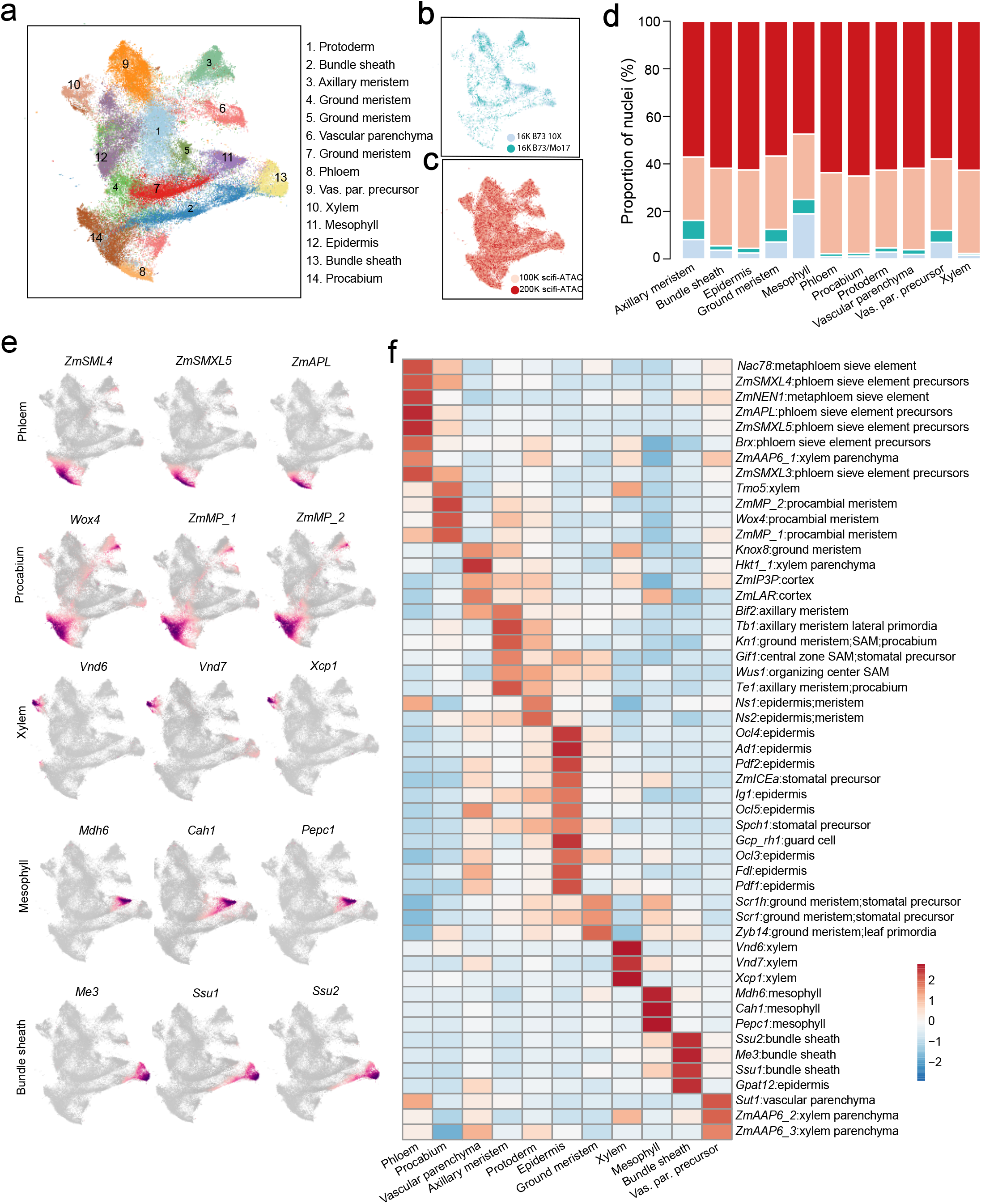
Clustering and cell-type annotation of scifi-ATAC-seq and scATAC-seq, related to Figure 1. (a) UMAP of all nuclei (n = 98,424). Nuclei are colored by clusters and labeled with predicted cell types. Vas. par. precursor: Vascular parenchyma precursor. (b) UMAP of nuclei from the two scATAC-seq dataset. Nuclei are colored by the library. (c) UMAP of nuclei from the two scifi-ATAC-seq datasets. Nuclei are colored by the library. (d) Proportions of nuclei derived from the four datasets for each cell type. Color scheme is the same with b and c. There was no significant bias in cell-type proportions between the two technologies (Spearman correlation: 0.62, p-value < 0.05). (e) UMAP embeddings overlaid with gene chromatin accessibility for representative marker genes. (f) Relative chromatin accessibility for the marker genes across all cell types.

**Figure S4.**
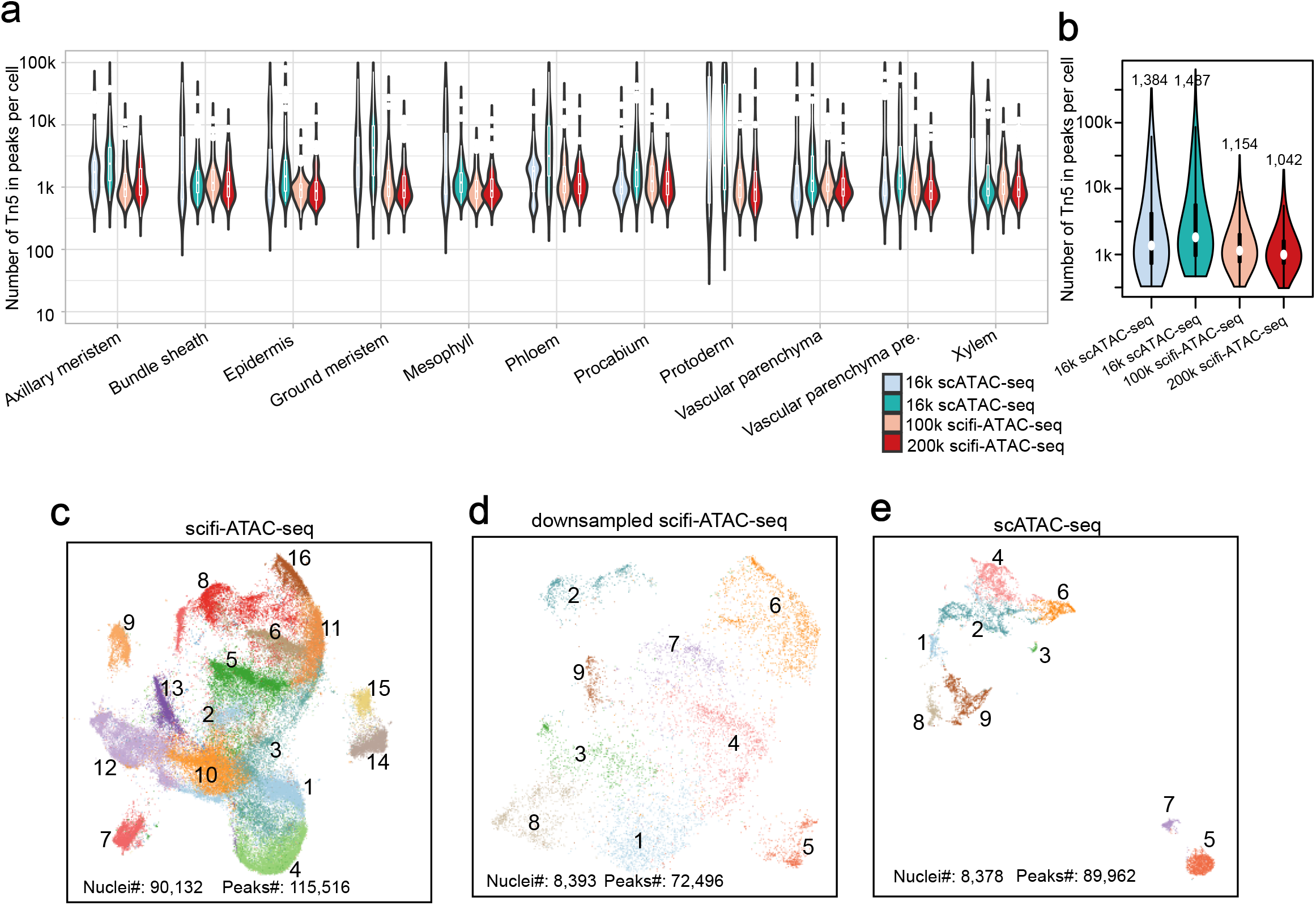
Comparison of nuclei quality between scifi-ATAC-seq and scATAC-seq, related to Figure 1. (a) Distribution of number of Tn5 integration for radomly selected 100 cells in each cell type and each library, summarized by cell type and library. (b) Distribution of number of Tn5 integration for radomly selected 100 cells in each cell type and each library, summarized by library. Median numbers were labeled for each library. (c) UMAP of all nuclei from the two scifi-ATAC-seq libraries. Nuclei are colored by clusters. (d) UMAP of downsampled nuclei from scifi-ATAC-seq. Nuclei are colored by clusters. (e) UMAP of all nuclei from the two scATAC-seq libraries. Nuclei are colored by clusters.

**Figure S5.**
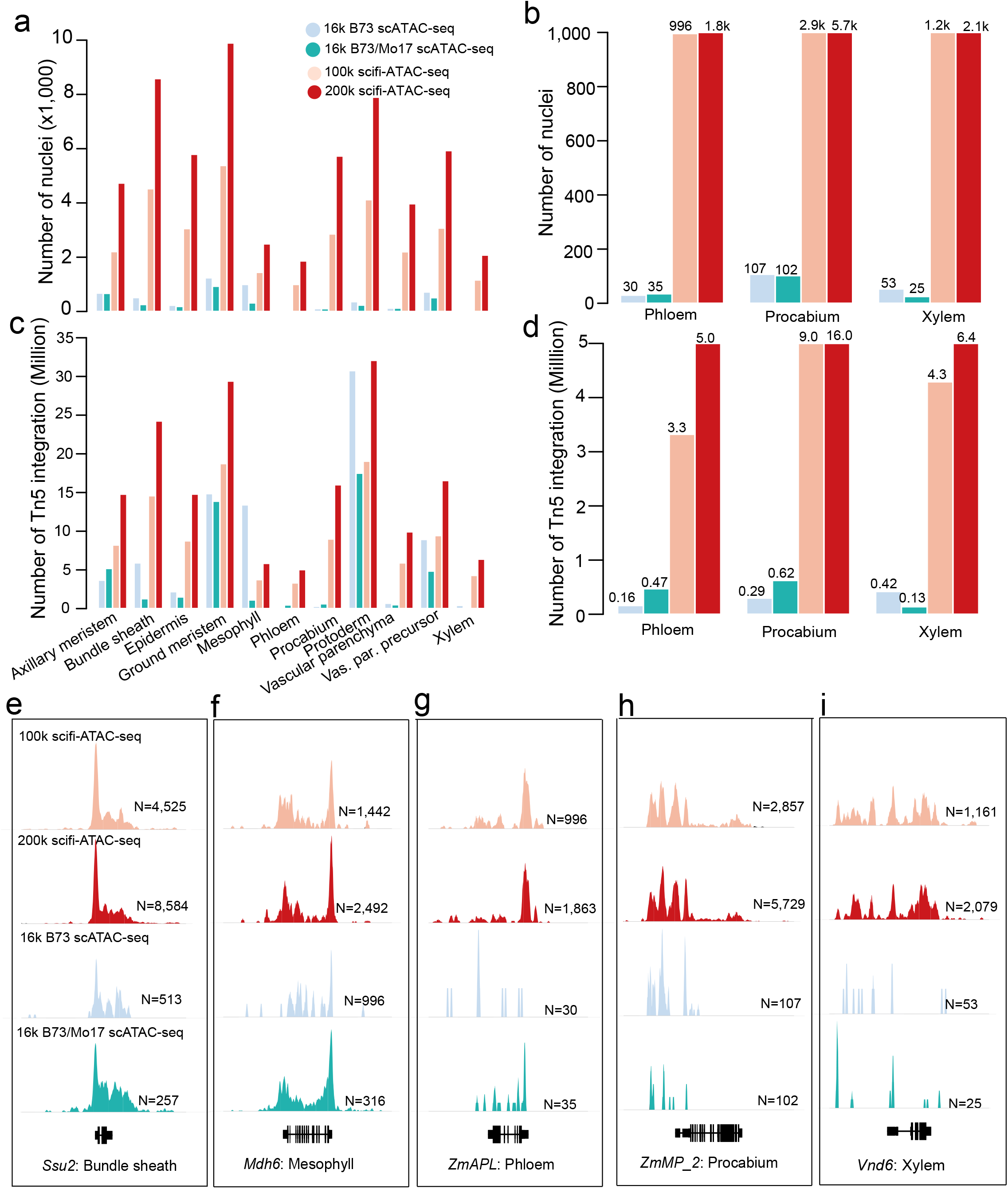
Characterization of vascular cells with scifi-ATAC-seq, related to Figure 1. (a) Overall distribution of nuclei number derived from each dataset across all cell types. (b) Distributions of nuclei number derived from each dataset across vascular cells, e.g. Phloem, Procambium and Xylem. The total nuclei number was labeled on the top of each bar. (c) Overall distribution of aggregated Tn5 integration numbers derived from each dataset across all cell types. (d) Distribution of aggregated Tn5 integration number derived from each dataset across vascular cells, e.g. Phloem, Procambium and Xylem. The total Tn5 integration numbers were labeled on the top of each bar. (e-i) Pseudobulk cell-type Tn5 integration site coverage at representative marker genes for vascular cells across four datasets. e, Bundle sheath; f, Mesophyll, g, Phloem; h, Procambium, i. Xylem. N: The total nuclei number according to cell type.

**Figure S6.**
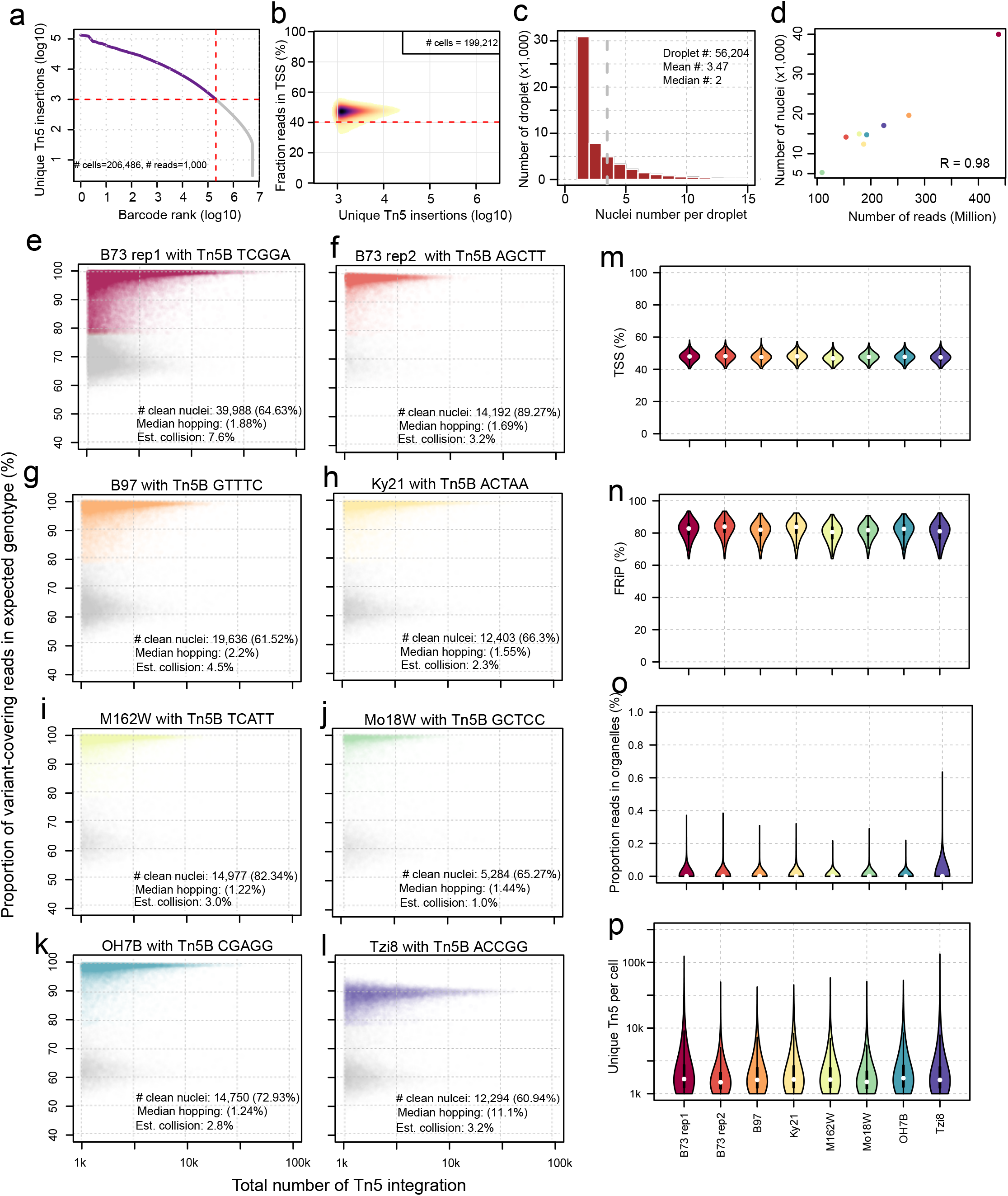
Evaluation and quality control of multiplexed scifi-ATAC-seq, related to Figure 2. (a) Identification of high-quality barcodes using a knee-plot. (b) Density scatterplots of log10 transformed barcode read depths (x axis) by the fraction of Tn5 integration sites mapping to within 2-kb of transcription start sites (TSSs). Dashed red lines indicate the threshold of two standard deviations from the mean used to filter lower quality barcodes. (c) Distributions of nuclei number per droplet in 300K input multiplexed scifi-ATAC-seq. (d) Correlation between sequencing depth and identified nuclei number for each library. (e-l) Distributions of proportion of variant-covering reads in expected genotypes matching known tn5 barcode for eight samples. Colored dots: clean nuclei. Grey dots: identified cell with high index hopping contamination. Median Contamination Rate: The median cross-contamination rate, attributed to index hopping, among all clean nuclei. (m) Distributions of the proportion of Tn5 integration sites within the promoter regions, encompassing the 2-kb flanking regions around gene transcription start sites (TSSs). (n) Distributions of the proportion of Tn5 integration sites within peaks per nucleus. (o) Distributions of the proportion of Tn5 integration sites within organelle genomes. (p) Distributions of unique Tn5 integration sites per nucleus.

**Figure S7.**
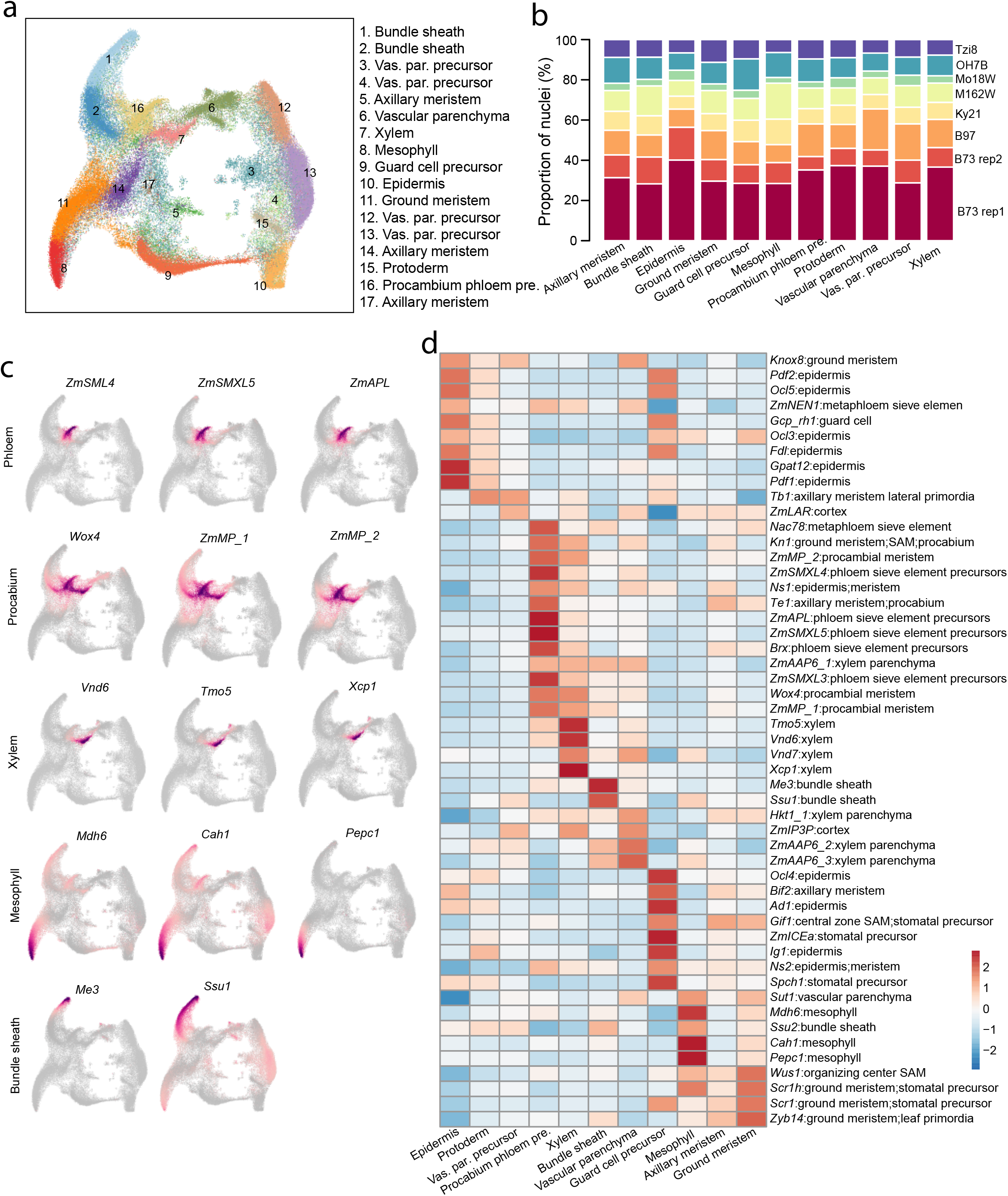
Clustering and cell-type annotation of multiplexed scifi-ATAC-seq, related to Figure 2. (a) UMAP of all nuclei (n = 124,656). Nuclei are colored by clusters and labeled with predicted cell types. Vas. par. precursor: Vascular parenchyma precursor. Procambium phloem pre.: procambium phloem precursor. (b) Proportion of nuclei derived from the eight samples for each cell type. (c) UMAP embeddings overlaid with gene chromatin accessibility for representative marker genes. (d) Relative chromatin accessibility for the marker genes across all cell types.

**Figure S8.**
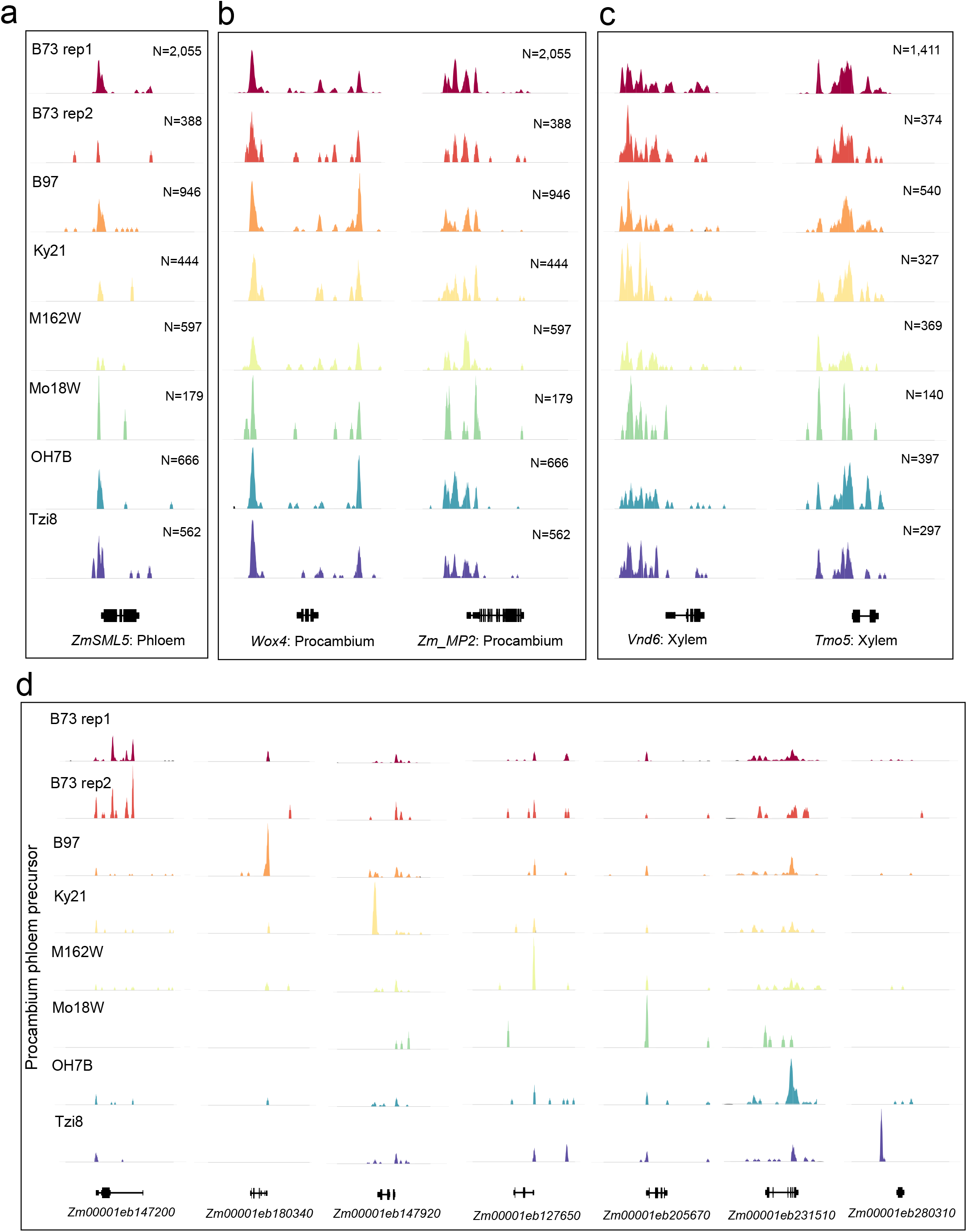
Characterization of vascular cells and genotype-specific accessible chromatin regions with demultiplexed scifi-ATAC-seq, related to Figure 2. (a-c) Pseudobulk cell-type Tn5 integration site coverage at representative marker genes for vascular cells across all samples. a, Phloem; b, Procambium, c. Xylem. N: The total nuclei number according to cell type. (e) Pseudobulk cell-type Tn5 integration site coverage at representative genotype-specific accessible chromatin regions in procambium phloem precursor cells.

**Figure S9.**
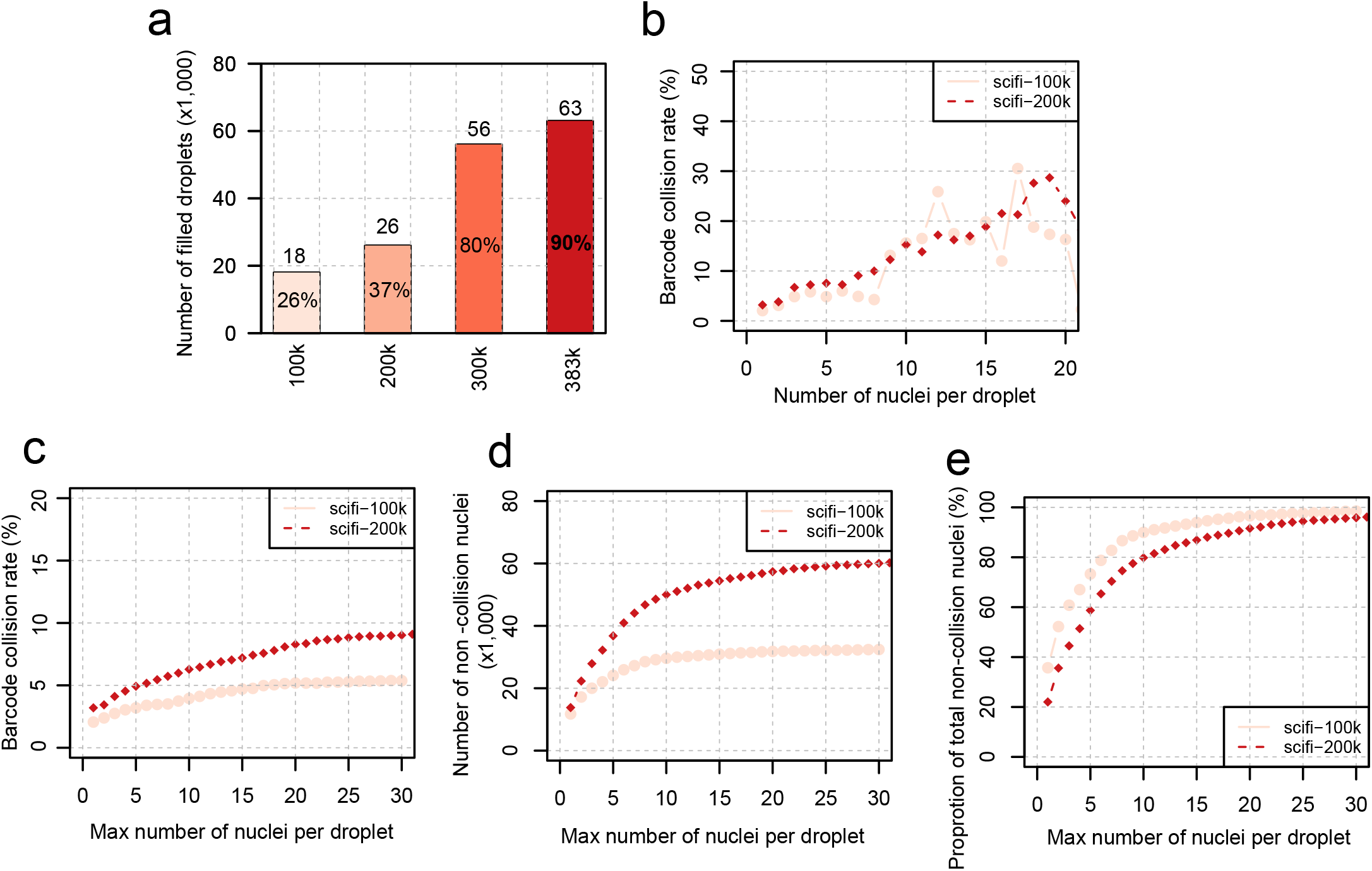
Estimation of barcode collision rate with different input nuclei for scifi-ATAC-seq. (a) Distribution of number of filled droplets with different input amount of nuclei. The 383k nuclei input were from scifi-RNA-seq[14] Estimated proportions of filled droplets were labeled in the center of the bars, assuming 70k droplets were generated in each run. (b) Distribution of identified barcode collision rates for droplets with nuclei numbers ranging from 1 to 20. (c) Distribution of identified barcode collision rates for different cutoffs of maximum nuclei numbers in each droplet, ranging from 1 to 30. (d) Distribution of number of non-collsion nuclei for different cutoffs of maximum nuclei numbers in each droplet, ranging from 1 to 30. (e) Distribution of the proportion of total identified non-collision nuclei for different cutoffs of maximum nuclei numbers in each droplet, ranging from 1 to 30.

## References

1. Buenrostro, J.D., et al., Transposition of native chromatin for fast and sensitive epigenomic profiling of open chromatin, DNA-binding proteins and nucleosome position. Nat Methods, 2013. 10(12): p. 1213–8.

2. Cusanovich, D.A., et al., A Single-Cell Atlas of In Vivo Mammalian Chromatin Accessibility. Cell, 2018. 174(5): p. 1309–1324 e18.

3. Domcke, S., et al., A human cell atlas of fetal chromatin accessibility. Science, 2020. 370(6518).

4. Marand, A.P., et al., A cis-regulatory atlas in maize at single-cell resolution. Cell, 2021. 184(11): p. 3041–3055 e21.

5. Zhang, K., et al., A single-cell atlas of chromatin accessibility in the human genome. Cell, 2021. 184(24): p. 5985–6001 e19.

6. Farmer, A., et al., Single-nucleus RNA and ATAC sequencing reveals the impact of chromatin accessibility on gene expression in Arabidopsis roots at the single-cell level. Mol Plant, 2021. 14(3): p. 372–383.

7. Dorrity, M.W., et al., The regulatory landscape of Arabidopsis thaliana roots at single-cell resolution. Nat Commun, 2021. 12(1): p. 3334.

8. Marand, A.P. and R.J. Schmitz, Single-cell analysis of cis-regulatory elements. Curr Opin Plant Biol, 2022. 65: p. 102094.

9. Satpathy, A.T., et al., Massively parallel single-cell chromatin landscapes of human immune cell development and intratumoral T cell exhaustion. Nat Biotechnol, 2019. 37(8): p. 925–936.

10. Cusanovich, D.A., et al., Multiplex single cell profiling of chromatin accessibility by combinatorial cellular indexing. Science, 2015. 348(6237): p. 910–4.

11. Vitak, S.A., et al., Sequencing thousands of single-cell genomes with combinatorial indexing. Nat Methods, 2017. 14(3): p. 302–308.

12. Ding, J., et al., Systematic comparison of single-cell and single-nucleus RNA-sequencing methods. Nat Biotechnol, 2020. 38(6): p. 737–746.

13. Lareau, C.A., et al., Droplet-based combinatorial indexing for massive-scale single-cell chromatin accessibility. Nat Biotechnol, 2019. 37(8): p. 916–924.

14. Datlinger, P., et al., Ultra-high-throughput single-cell RNA sequencing and perturbation screening with combinatorial fluidic indexing. Nat Methods, 2021. 18(6): p. 635–642.

15. Tu, X., et al., A combinatorial indexing strategy for low-cost epigenomic profiling of plant single cells. Plant Commun, 2022. 3(4): p. 100308.

16. Granja, J.M., et al., ArchR is a scalable software package for integrative single-cell chromatin accessibility analysis. Nat Genet, 2021. 53(3): p. 403–411.

17. Germain, P.L., et al., Doublet identification in single-cell sequencing data using scDblFinder. F1000Res, 2021. 10: p. 979.

18. Thibodeau, A., et al., AMULET: a novel read count-based method for effective multiplet detection from single nucleus ATAC-seq data. Genome Biol, 2021. 22(1): p. 252.

19. Xi, N.M. and J.J. Li, Benchmarking Computational Doublet-Detection Methods for Single-Cell RNA Sequencing Data. Cell Syst, 2021. 12(2): p. 176–194 e6.

20. Tu, X., et al., Reconstructing the maize leaf regulatory network using ChIP-seq data of 104 transcription factors. Nat Commun, 2020. 11(1): p. 5089.

21. Lu, Z., et al., Combining ATAC-seq with nuclei sorting for discovery of cis-regulatory regions in plant genomes. Nucleic Acids Res, 2017. 45(6): p. e41.

22. Sikorskaite, S., et al., Protocol: Optimised methodology for isolation of nuclei from leaves of species in the Solanaceae and Rosaceae families. Plant Methods, 2013. 9: p. 31.

23. Smith, T., A. Heger, and I. Sudbery, UMI-tools: modeling sequencing errors in Unique Molecular Identifiers to improve quantification accuracy. Genome Res, 2017. 27(3): p. 491–499.

24. Martin, M., Cutadapt removes adapter sequences from high-throughput sequencing reads. EMBnet. journal, 2011. 17(1): p. 10–12.

25. Hufford, M.B., et al., De novo assembly, annotation, and comparative analysis of 26 diverse maize genomes. Science, 2021. 373(6555): p. 655–662.

26. Li, H. and R. Durbin, Fast and accurate short read alignment with Burrows-Wheeler transform. Bioinformatics, 2009. 25(14): p. 1754–60.

27. Danecek, P., et al., Twelve years of SAMtools and BCFtools. Gigascience, 2021. 10(2).

28. Canaran, P., et al., Panzea: an update on new content and features. Nucleic Acids Res, 2008. 36(Database issue): p. D1041–3.

29. Zhao, H., et al., CrossMap: a versatile tool for coordinate conversion between genome assemblies. Bioinformatics, 2014. 30(7): p. 1006–7.

30. Heaton, H., et al., Souporcell: robust clustering of single-cell RNA-seq data by genotype without reference genotypes. Nat Methods, 2020. 17(6): p. 615–620.

31. Korotkevich, G., et al., Fast gene set enrichment analysis. BioRxiv, 2016: p. 060012.

32. Zhang, Y., et al., Model-based analysis of ChIP-Seq (MACS). Genome Biol, 2008. 9(9): p. R137.

33. Robinson, M.D., D.J. McCarthy, and G.K. Smyth, edgeR: a Bioconductor package for differential expression analysis of digital gene expression data. Bioinformatics, 2010. 26(1): p. 139–40.

34. Zhang, Hao, et al. txci-ATAC-seq, a massive-scale single-cell technique to profile chromatin accessibility. bioRxiv (2023): 2023–05.

35. Yan, Haidong, et al. Evolution of cell-type-specific accessible chromatin regions and the cis-regulatory elements that drive lineage-specific innovation. bioRxiv (2024): 2024–01.

36. Mendieta, John Pablo, et al. Investigating the cis-Regulatory Basis of C3 and C4 Photosynthesis in Grasses at Single-Cell Resolution. bioRxiv (2024): 2024–01.

