## Supplementary Method scifi-ATAC-seq protocol for "Massive-scale single-cell chromatin accessibility sequencing using combinatorial fluidic indexing"

### Supplementary Methods

**Scifi-ATAC-seq protocol.** Here, we report scifi-ATAC-seq. The developed approach should be applicable to a wide range of species, we have also applied it to soybean successfully (data not shown). A step-by-step protocol is presented below in detail.

#### I. Adapter and primer design:

1) Tn5-ME-A index adapter A1-A12 (Supplementary Table S1):

```
5′-TCGTCGGCAGCGTCGATATGTGATAATGAGGAC NNNNN AGATGTGTATAAGAGACAG-3′
                                        3′-TCTACACATATTCTCTGTC-phos-5′
```

2) Tn5-ME-B index adapter B1-B8 (Supplementary Table S1):

```
5′-GTCTCGTGGGCTCGGTGAATGTGTAGAAGACAGA NNNNN AGATGTGTATAAGAGACAG-3′
                                         3′-TCTACACATATTCTCTGTC-phos-5′
```

3) qPCR primer set (10 µM):

```
scifi-qPCR-F: 5'-TCGTCGGCAGCGTCGATATGTGATAATGAGGAC-3'
scifi-qPCR-R: 5'-GTCTCGTGGGCTCGGTGAATGTGTAGAAGACAGA-3'
```

4) Custom sequencing primers set (HPLC purified, 100 µM):

```
1_Read1: 5′-TCGTCGGCAGCGTCGATATGTGATAATGAGGAC-3′
2_Index1(i7): 5′-TCTGTCTTCTACACATTCACCGAGCCCACGAGAC-3′
3_Index2(i5): 5′-GTCCTCATTATCACATATCGACGCTGCCGACGA-3′
4_Read2: 5′-GTCTCGTGGGCTCGGTGAATGTGTAGAAGACAGA-3′
```

#### II. Anneal index Tn5 adapters

1. Dissolve 12 Tn5-ME-Ax and 18 Tn5-ME-Bx with the complementary Tn5-ME-Rev single stranded DNA oligonucleotides (oligos) with TE to 200 µM.
2. Mix equal volume of Tn5-ME-Ax and Tn5-ME-Rev in a PCR tube to obtain a 100 µM adapter stock.
3. Similarly, mix equal volume of Tn5-ME-Bx and Tn5-ME-Rev.
4. Heat the oligo mixture to 98°C in a PCR instrument for 2 min. and gradually lower the temperature to 25°C by cycling (-1°C per 10 sec cycle).
5. The annealed adapters can be used immediately or stored at -20 °C.

#### III. Assemble 96 indexed TS-Tn5

1. To assemble Tn5 with A1-A12 adapters, mix 2 µL annealed Ax adapter with 15 µL Tn5 in PCR 8-well strip tubes. Mix well by pipetting about 20 times slowly and avoid generating bubbles.
2. Similarly, to assemble Tn5 with B1-B8 adapters, mix 3 µL Bx adapter with 25 µL diluted Tn5 in PCR 8-well strip tube.
3. Incubate the PCR strips at 25 °C for 1 h to assemble the Tn5.
4. The assembled Tn5 can be used immediately, or stored at -20 °C.
5. Distribute 1.5 µL Tn5 with adapter B1-B8 to each column of the PCR 96-well reaction plate on ice.
6. Similarly, distribute 1.5 µL Tn5 with adapter A1-12 to each row of the PCR 96 well reaction plate on ice.
7. Each well now contains 3 µL of well assembled Tn5 with distinct index combinations.
8. The plates can be used immediately or sealed for overnight temporary storage at -20 °C.

**IV. Nuclei isolation with quick purification**

1. Prepare 10 mL 4x **Nuclei Isolation Buffer** (4x NIB) on ice, can store at 4 °C up to 1 month.

4x Nuclei Isolation Buffer(4x NIB):

| Stock | Final(4x) | 10 mL |
|---|---|---|
| 0.4M MES-KOH pH 5.4 | 40 mM | 1 mL |
| 5M NaCl | 40 mM | 40 µL |
| 2M sucrose | 1 M | 5 mL |
| $H_2O$ | | to 10 mL |

2. Prepare 10 mL **NIB-wash buffer** on ice:

| Stock | Final(1x) | 10 mL |
|---|---|---|
| 4x NIB | 1x | 2.5 mL |
| 1M Spermine | 0.1 mM | 1 µL |
| 1M Spermidine | 0.5 mM | 5 µL |
| 1M DTT | 1 mM | 10 µL |
| 10% BSA | 1% | 1 mL |
| $H_2O$ | | to 10 mL |

3. Prepare **2x TAPS buffer**, can store at -20 °C:

| Stock | Final(2x) | 100 mL |
|---|---|---|
| TAPS(Alfa Aesar, cat#: A17754) | 50 mM | 1.22g, dessove in 80 mL $H_2O$, adjust pH to 8.0 with NaOH |
| 1M $MgCl_2$ | 25 mM | 2.5 mL |
| $H_2O$ | | to 100 mL |

4. Mix 5 mL NIB-wash buffer and 250 µL 10% TritonX-100 to make the NIB-cutting buffer.
5. Approximately 3-4 maize seedlings (7-10 day old) were placed on a petri dish on ice and saturated with 500 mL of chilled **NIB-cutting buffer**.
6. Using a sterilized single-edge razor blade, seedlings were chopped for about 2 min. to break the cell wall, lyse the chloroplasts and mitochondria, and release nuclei into solution.
7. The aqueous nuclei slurry was then filtered through a 40 µm cell strainer (pluriSelect, cat# 43-10040-40).
8. Pellet the nuclei by centrifugation (swinging-bucket centrifuge rotor) at 500 rcf for 5 min. at 4°C.
9. Remove supernatant carefully and leave some of the supernatant behind if necessary.
10. Resuspend the nuclei pellet gently with 500 µL **NIB-wash buffer**, filter through a 20 µm cell strainer (pluriSelect, cat# 43-10020-60).
11. Slowly add the nuclei suspension on the top of 1 mL 35% percoll buffer (mixed 350 µL Percoll and 650 µL NIB wash buffer).
12. Centrifuge at 500 rcf for 10 min. at 4°C. This step further removes the chloroplasts, mitochondria, and small debris.
13. Slowly remove supernatant from the top to the bottom, and leave about 10 µL of supernatant to avoid removing any nuclei.
14. Resuspend nuclei with about 200 µL TAPS buffer (with 0.1% Tween 20 and 0.01% digitonin), centrifuge 500 rcf for 2 min.
15. Remove the supernatant and resuspend in 30 µL TAPS buffer.
16. Take 5 µL nuclei to a centrifuge tube, dilute 10 fold with TAPS buffer, add 0.2 µL DAPI(1 mg/mL). Load 5 µL diluted nuclei to a hemocytometer and check the nuclei quality and density under microscope.

17. Dilute the nuclei density to about 0.5-1k/µL with TAPS buffer(with 0.1% Tween 20 and 0.01% digitonin).

**V. Indexed Tn5 tagmentation and pooling.**

1. Distribute 10 µL nuclei to each well of the PCR 96 well plate and mix with indexed Tn5 from the previous procedure by pipetting 10 times slowly.
2. Incubate the tagmentation plate at 37°C for 60 min.
3. After Tn5 treatment, add 12 µL stop buffer each well, Mix well by pipetting 5 times slowly.

| Stock | Final(1x) | 4 mL |
|---|---|---|
| 1M Tris-HCl (pH 7.8) | 10 mM | 40 µL |
| 0.5M EDTA | 20 mM | 160 µL |
| 10% BSA | 2% | 800 µL |
| $H_2O$ | | to 4 mL |

4. Pool all nuclei together and transfer it to two 1.5 mL centrifuge tubes. Pellet the nuclei at 500 rcf for 10 min. at 4°C, aspirate supernatant and resuspend in 200 µL diluted nuclei buffer (DNB, 10x Genomics Cat#2000207)
5. Then filter the nuclei with a 40 µm cell strainer, check the nuclei quality and density, and transfer 200k-400k nuclei to a new PCR tube.
6. Pellet the nuclei at 500 rcf for 2 min at 4°C.
7. The supernatant was carefully removed, leave about 5 µL of buffer in the tube, and mix with 7 µL of ATAC buffer B (10x Genomics Cat#2000193).

**VI. Library preparation and sequencing**

1. Follow manufacturer's instruction ( 10xGenomics, CG000209_Chromium_NextGEM_SingleCell_ATAC_ReagentKits_v1.1_UserGuide_RevE) exactly for the Gel Beads-in-emulsion (GEM) generation & barcoding and post GEM incubation cleanup. (Note: To estimate accurate input nuclei number, we also dilute the leftover nuclei 5 times after loading to the ChIP-H, stain with DAPI, and check nuclei quality and density with a hemocytometer under microscope.)

2. With the aim of determining PCR cycles prior to saturation occurs to mitigate amplification bias for the library construction, 1 µL of DNA sample from above was spared to perform the 10 µL real-time PCR (qPCR):

    5 µL of 2x Luna® Universal qPCR Master Mix(NEB, cat# M3003 )
    0.5 µL Scifi-qPCR-F
    0.5 µL Scifi-qPCR-R
    1 µL DNA Sample
    3 µL Nuclease-Free Water

3. Program Bio-Rad CFX96 qPCR system and perform the thermal profile as below:

| | |
|---|---|
| Initial denaturation for 30s at 95°C | |
| | Denature for 15s at 95°C |
| 30 cycles | Anneal primers for 30s at 63°C |
| Detect fluorescence | Extend DNA for 1 min at 72°C |
| 4°C Hold | |

4. Following the library construction from the manufacturer's instruction, Amplify the remaining 40 µL sample using a suitable cycle number before saturation based on the qPCR test. (Note we usually amply the cycle number around Cq value, and we aim to obtain ~ 100 ng library DNA without more than 9 additional cycles.)
5. The final libraries were sequenced with Illumina NovaSeq 6000 S4 in dual-index mode with the custom sequencing primer sets. The final library was pooled with libraries from an unrelated experiment to balance the nucleotide distribution at the beginning of the forward and reverse reads. Usually, we keep the proportion of the scifi-ATAC-seq library in a lane to be less than 50%, or more spike-in library (e.g.,PhiX control from Illumina) should be added to the lane.

#### Supplementary Note:

1. Application of scifi-ATAC-seq for different species and tissues:

In theory, there should be no sample or species limits for scifi-ATAC-seq, provided that high-quality nuclei can be prepared. We have successfully obtained high-quality nuclei for numerous plant species with the current nuclei isolation protocol, including maize, soybean, rice, wheat, barley, Arabidopsis, and poplar, across various tissue types such as leaf, root, nodule, hypocotyl, and developing seed. We recommend validating this nuclei isolation protocol by assessing nuclei quality under a microscope and evaluating the data quality through regular/non-single-cell ATAC-seq prepared from the isolated nuclei.

2. Maximum samples in one scifi-ATAC-seq assay:

In theory, it is feasible to mix up to 96 samples in one assay, given the use of 96 barcodes in this study. However, in practice, there is a risk of nuclei degradation if they remain in the buffer for an extended period. The nuclei isolation for each sample takes approximately 4 minutes, and an additional 30 minutes are needed for nuclei purification for all samples. In the 8-sample scifi-ATAC-seq, we aim to complete the entire nuclei preparation within 1 hour to minimize the risk of nuclei degradation during the process. Including more samples is possible if a sufficient number of high-quality nuclei can be obtained in the given time frame. From the 8-sample scifi-ATAC-seq with a 300k input, we obtained about 120k nuclei following strict QC. Therefore, it should be feasible to include up to 12 samples in the 300k input scifi-ATAC-seq assay, ensuring an average of 10k nuclei per sample, which is generally acceptable for most experiments. In animals, a similar approach called txci-ATAC-seq has been developed. It provides a protocol for preparing scATAC-seq with frozen nuclei, offering the potential to profile up to 96 samples simultaneously [34].

3. Maximum numbers of nuclei in one scifi-ATAC-seq assay:

The maximum number of nuclei is primarily determined by the number of unique barcodes, with more indexes reducing the probability of barcode collision when multiple nuclei are present in one droplet. In other words, in the 96-barcodes experiment, the probability of barcode collision in the droplet with a certain number of nuclei is expected to be similar, regardless of the total number of loaded nuclei. To verify this, we compared barcode collision rates for droplets containing one to twenty nuclei in both 100k and 200k input scifiATAC-seq. The results showed an increase in the barcode collision rate with more nuclei in the droplet, yet showing similarity between 100k and 200k input scifi-ATAC-seq, where the collision rate is approximately 15% for droplets with 10 nuclei (Fig S9b).

However, the distribution of the number of nuclei in droplets for 100k, 200k, and 300k input nuclei is not a normal distribution but a biased distribution, with most droplets containing one to ten nuclei (Fig S1d,f; Fig S6c). Therefore, it is possible to further decrease the barcode collision rate by removing droplets with a high number of nuclei. For instance, the barcode collision rate could be reduced to about 4% and 6% when filtering droplets with more than 10 nuclei while retaining over 90% and 80% of the total non-collision nuclei for 100k and 200k input scifi-ATAC-seq, respectively (Fig S9c-e).

In one assay from the 10X Genomics Chromium Controller, approximately 70k droplets were generated, and about 90% of these droplets could be filled with 383k input nuclei (Fig S9a). However, as the number of input nuclei increases, there is a higher risk of chip clogging due to the high density of nuclei. We believe it might be feasible to load 400k nuclei in each run and selectively retain only the droplets with no more than 10 nuclei for downstream analysis using the 96-barcodes scifi-ATAC-seq method.

4. Potential limits for mixing samples from different species in one assay:

There is a potential for sequencing throughput bias among nuclei from different species if they contain significantly different sizes of chromatin accessibility regions due to variations in genome size and as well as GC content variation among species.
